# Serotonergic Modulation of Motoneuronal Excitability and Its Impact on Muscle Force Generation: A Computational Study

**DOI:** 10.1101/2025.04.01.646607

**Authors:** José Meléndez-Gallardo, Dinorah Plada Delgado

## Abstract

This study investigates the impact of serotonin (5-HT) on motoneuron electrical activity and muscle force generation. Using a computational model, we explore how 5-HT receptors influence motoneuron excitability and muscle function at different stimulation frequencies. Our results demonstrate that physiological 5-HT release increases motoneuron excitability, particularly at higher frequencies (40 Hz and 100 Hz), consistent with the known excitatory role of serotonin through 5-HT2a receptor activation. However, high concentrations of 5-HT lead to decreased motoneuron excitability, potentially due to excessive activation of 5-HT1a receptors, which are involved in neuronal hyperpolarization. The generated muscle force exhibited a direct correlation with motoneuron activity. At 10 Hz, physiological 5-HT release did not significantly affect force production, suggesting minimal impact at low frequencies. However, high 5-HT concentrations reduced contraction duration, indicating muscle fatigue. At 40 Hz and 100 Hz, physiological 5-HT release enhanced motoneuron excitability, promoting more sustained muscle responses. Nevertheless, high serotonin concentrations decreased the ability to maintain muscle contraction, leading to fatigue. Our findings align with experimental studies, validating the model as an effective tool for understanding the role of serotonin in neuromuscular function. The model provides insights into the mechanisms underlying muscle fatigue and could serve as a foundation for developing therapeutic strategies to modulate serotonergic signaling, improving muscle performance and recovery. Future research should explore the interactions with other neurotransmitters to further investigate their effects on motor function and fatigue resistance.

## Introduction

The function of skeletal muscle is determined by the activation of motor units (MUs), which are modulated by motoneurons in response to synaptic signals. In this process, neurotransmitters play a key role in motoneuron excitability and, consequently, in muscle force production [1]. Among these neurotransmitters, serotonin (5-HT) is particularly relevant, as its action on different receptors can modulate muscle fatigue and motor control efficiency [2].

However, the interaction between 5-HT1a and 5-HT2a receptors in neuronal excitability and their impact on force production remains poorly understood. It has been demonstrated that 5-HT1a and 5-HT2a receptors differentially affect motoneuron excitability [3]. While the 5-HT1a receptor inhibits neuronal activity and contributes to fatigue, the 5-HT2a receptor promotes sustained motoneuron activation, suggesting an adaptive mechanism to maintain muscular effort without a rapid decline in performance. Nevertheless, an imbalance in the activation of these receptors can be detrimental, as elevated serotonin levels have been linked to central fatigue [4].

During muscle activity, neurons in the raphe nucleus release 5-HT onto the dendrites and soma of motoneurons, where excitatory 5-HT2a receptors are abundant. However, if neurotransmitter release is excessively high, inhibitory 5-HT1a receptors are also activated. These receptors are located in the initial segment of the axon and modulate action potential propagation by inhibiting Na+ channels [5].

To better understand the role of serotonin in muscle fatigue modulation, it is crucial to develop computational models that accurately replicate the interaction between serotonergic receptors and motoneurons. This study presents a model using Python’s NEURON module, simulating motoneuron activation in response to serotonin release from a presynaptic neuron. The motoneuron is further connected to a muscle model, enabling the evaluation of force production as a function of excitability modulated by 5-HT. This approach allows for the exploration of how differential activation of 5-HT1a and 5-HT2a receptors influences muscle contraction dynamics and fatigue generation.

## Materials and Methods

### Technical information

The model consists of a motoneuron and a generic supraspinal presynaptic neuron (raphe nuclei) that releases neurotransmitter 5-HT upon stimulation [5]. The modeling was developed with the Neuron 8.2 module of Python 3.12, in the Spyder development environment. This module is a simulation environment for modeling individual and network neurons, which provides a range of conventional tools for constructing, managing, and using modeling in a numerically robust and computationally efficient manner [6]. The scripts files of the model developed in this study are publicly available at: https://github.com/JGMG7/Serotonergic-Modulation-of-Motoneuronal-Excitability-and-Muscle-Force-Generation-Computational-Study

### Motoneuron

The motoneuron models used in this study were based on the works of Kim (2017) [7], Fietkiewicz et al. (2023) [6], and Meléndez-Gallardo (2024) [8]. This model consists of a multi-compartmental cable model, with anatomical data corresponding to a cat motoneuron (i.e., v_e_moto6), which is available in the public database (www.neuromorph.org). The non-uniform specific membrane resistivity was assigned to the soma (Rm, soma = 225 Ω·cm^2^) and the dendrites (Rm, dendrite = 225 Ω·cm^2^). The specific membrane capacitance (Cm = 1 μF/cm^2^) and axial resistivity (Ri = 70 Ω·cm) were uniformly assigned to all compartments of the motoneuron model. Action potentials and afterhyperpolarization phenomena were generated by various Hodgkin-Huxley-type active currents: a fast-inactivating sodium current (I_Naf), a delayed rectifier potassium current (I_KDr), a persistent sodium current (I_Nap), an N-type calcium current (I_CaN), and a calcium-dependent potassium current [I_K(Ca)] in the soma. Additionally, I_Naf, I_Nap, and I_KDr were included in the initial segment/axon hillock. The peak conductances for active currents were G_Naf = 0.71 S/cm^2^, G_KDr = 0.23 S/cm^2^, G_CaN = 0.013 S/cm^2^, and G_K(Ca) = 0.0258 S/cm^2^ at the soma, while at the initial segment/axon hillock, they were G_Naf = 2.7 S/cm^2^, G_Nap = 0.033 mS/cm^2^, and G_KDr = 0.17 S/cm^2^. To generate the inhibitory and excitatory response of the motoneuron, 5-HT receptors of subtypes 5-HT1a and 5-HT2a were incorporated into the initial segment, dendrites, and soma, respectively [3].

### Muscle unit

The muscle unit consists of a section that incorporates two mechanisms: a calcium mechanism and a force mechanism. The force mechanism relies on variables from the calcium mechanism for its activation. Additionally, the muscle communicates with the motoneuron’s spike timing through a NetCon connection, utilizing pointers to ensure smooth interaction between biomechanically distinct elements. The equations governing these processes are detailed in Fietkiewicz et al. (2023) [6].

### Presynaptic Neuron

The presynaptic neuron is modeled as a generic neuron (raphe nuclei) whose primary function is to release neurotransmitters upon receiving a stimulus (IClamp). It contains voltage-dependent Na^+^ channels (with three states) and K^+^ channels (with two states), following a Markov model [8,9].

It is assumed that upon the arrival of an action potential, the presynaptic terminal depolarizes, allowing Ca^++^ ions to enter, generating a high-threshold Ca^++^ current. These calcium ions then activate a calcium-binding protein, which facilitates the release of serotonin (5-HT) from presynaptic vesicles into the synaptic cleft. The presynaptic vesicles are considered inexhaustible and always available for release. This process is modeled as a first-order reaction with a stoichiometric coefficient of n.

The entry of calcium into the presynaptic terminal is governed by a high-threshold Ca ^++^ current, using the same two-state Markov scheme as the K^+^ channel, with a voltage-dependent rate similar to that of the K^+^ current. Intracellular Ca^++^ removal is facilitated by active transport via a calcium pump.

The 5-HT release mechanism interacts with 5-HT1a and 5-HT2a receptors on the motoneuron through the C pointer, which is, in turn, regulated by variables present in the rel2 mechanism (such as vesicle concentration, neurotransmitter molecules per vesicle, neurotransmitter hydrolysis rate, etc.) [8,9].

### 5-HT receptors

The 5-HT1a and 5-HT2a receptors were designed based on the kinetic response of G protein-coupled membrane receptors, which modulate ion channels [8,9]. The activity of both receptors is regulated by variables from the rel2 release mechanism of the presynaptic neuron via the C pointer. Specifically, for the 5-HT1a receptor, variations in the C pointer influence Ca^++^ currents in the motoneuron, while for the 5-HT2a receptor, fluctuations in the C pointer modulate Na+ and K^+^ currents. The association (ka) and dissociation (kd) constants used to model the 5-HT receptors were calculated as the average of the inhibition constant (ki) values reported in the literature [10].

Additionally, each receptor model includes the ability to inhibit 5-HT reuptake, leading to increased neurotransmitter availability in the extracellular space. This mechanism is based on modifications to the variable kh, which acts as a constant that regulates the 5-HT hydrolysis rate, modeled as a first-order reaction [8,9]. The 5-HT2a receptor was inserted into the soma and the 311 dendrites of the motoneuron, while the 5-HT1a receptor was located in the initial segment.

### Stimulation Protocols

The stimulation protocol consisted of a NetStim generating simultaneous input to both the soma and dendrites (311) of the motoneuron, as well as to the raphe nucleus neuron (presynaptic neuron). The synaptic characteristics were set as weight = 2, tau = 0.5, delay = 0, while the number of stimuli and interstimulus interval could be adjusted based on the required stimulus intensity.

### Neurotransmitter Release

To simulate 5-HT release, we manipulated the nt variable in the rel2 neurotransmitter release mechanism. The nt variable defines the number of molecules released each time the presynaptic neuron (raphe nucleus neuron) is excited. For what we defined as physiological release, the variable was set to nt = 10,000 molecules of 5-HT. For high-concentration neurotransmitter release, the variable was set to nt = 1,000,000.

### Statistical Analysis

The collected data were exported in.csv format and analyzed using the SciPy Python library [11] within the Spyder virtual environment. The Kruskal-Wallis test was applied, followed by Dunn’s multiple comparisons test. Significance threshold: p < 0.05 = significant, ns = not significant.

## Results

The motoneuron responses and muscle force generation were evaluated at three different stimulation frequencies (10 Hz, 40 Hz, and 100 Hz) under three distinct conditions, represented in the figures as a, b, and c: (a) No 5-HT receptor activity, (b) Physiological 5-HT release, (c) High-concentration 5-HT release. Muscle force recordings are also presented in the figures as d, e, and f, corresponding to conditions a, b, and c, respectively.

### Motoneuron Electrical Activity and Muscle Force Generation at 10 Hz Stimulation

Motoneuron electrical activity varied significantly depending on the presence of 5-HT receptors and the intensity of serotonergic discharge. At a stimulation frequency of 10 Hz: No significant differences were observed between the motoneuron activity without 5-HT receptors (condition a) and the motoneuron with 5-HT1a and 5-HT2a receptors under physiological 5-HT release (condition b) (p > 0.05). However, a high 5-HT discharge (condition c) resulted in a significant decrease in electrical activity compared to both conditions a and b (p < 0.0001 in both cases) (Table 1, Figure 1).

**Table 1.** Statistical Analysis of Motoneuron Electrical Activity (Vm) Under Different Stimulation Frequencies.

| **Stimuli 10 Hz** | **Difference in rank sum** | **p** | **Summary** |
| --- | --- | --- | --- |
| Motoneuron (a) vs Motoneuron (b) | 505,2 | >0.05 | ns |
| Motoneuron (a) vs Motoneuron (c) | 55860 | <0.0001 | *** |
| Motoneuron (b) vs Motoneuron (c) | 55350 | < 0.0001 | *** |
| **Stimuli 40 Hz** | **Difference in rank sum** | **p** | **Summary** |
| Motoneuron (a) vs Motoneuron (b) | 1661 | <0.0001 | *** |
| Motoneuron (a) vs Motoneuron (c) | 53690 | <0.0001 | *** |
| Motoneuron (b) vs Motoneuron (c) | 52020 | <0.0001 | *** |
| **Stimuli 100 Hz** | **Difference in rank sum** | **p** | **Summary** |
| Motoneuron (a) vs Motoneuron (b) | 1966 | <0.0001 | *** |
| Motoneuron (a) vs Motoneuron (c) | 40990 | <0.0001 | *** |
| Motoneuron (b) vs Motoneuron (c) | 39020 | <0.0001 | *** |

*Kruskal-Wallis Test and Dunn's Multiple Comparison Test. $p < 0.05$ = significant, ns = not significant*

**Figure 1.**
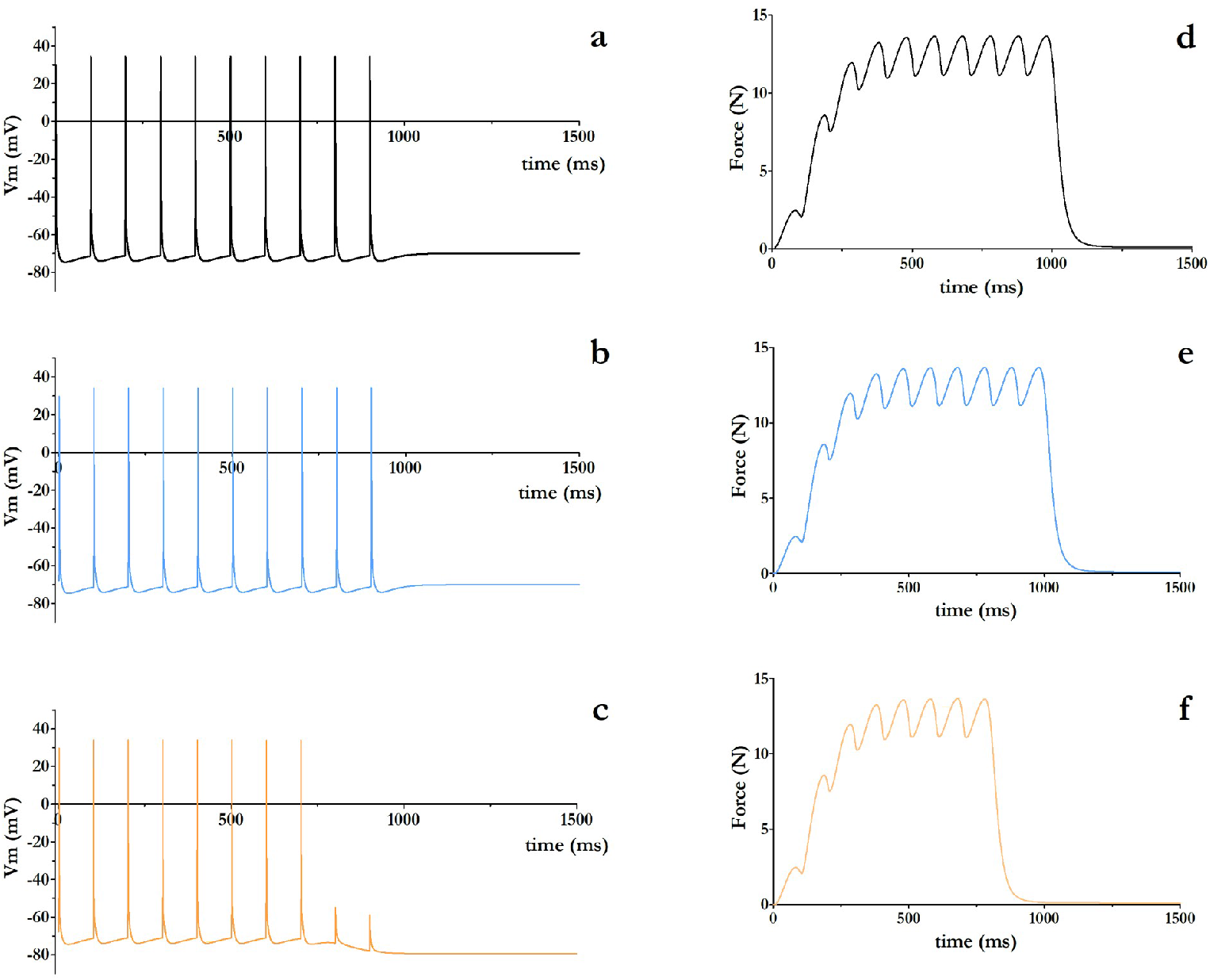
Motoneuron Membrane Potential and Muscle Force Generation at a Stimulation Frequency of 10 Hz. Motoneuron membrane potential without 5-HT receptors (a), with 5-HT1a and 5-HT2a receptors during physiological 5-HT release (b), and with 5-HT1a and 5-HT2a receptors during high 5-HT release (c).(d-f) Muscle force generated by the muscle innervated by the corresponding motoneuron in (a), (b), and (c), respectively.

In terms of muscle force, at 10 Hz, no significant differences were observed between conditions (a) and (b) (p > 0.05), indicating that physiological 5-HT release does not significantly alter muscle force generation at low stimulation frequencies. However, high-concentration 5-HT release (c) led to a reduction in force curve duration, suggesting the induction of fatigue (p < 0.0001) (Table 2, Figure 1).

**Table 2.** Statistical Analysis of Muscle Force (N) Generated Under Different Stimulation Frequencies.

| **Stimuli 10 Hz** | **Difference in rank sum** | **p** | **Summary** |
| --- | --- | --- | --- |
| Muscle Force (a) vs Muscle Force (b) | 0,0000 | >0.05 | ns |
| Muscle Force (a) vs Muscle Force (c) | 18530 | <0.0001 | *** |
| Muscle Force (b) vs Muscle Force (c) | 18530 | <0.0001 | *** |
| **Stimuli 40 Hz** | **Difference in rank sum** | **p** | **Summary** |
| Muscle Force (a) vs Muscle Force (b) | 0,0000 | >0.05 | ns |
| Muscle Force (a) vs Muscle Force (c) | 36850 | <0.0001 | *** |
| Muscle Force (b) vs Muscle Force (c) | 36850 | <0.0001 | *** |
| Muscle Force (a) vs Muscle Force (b) | 0,0000 | >0.05 | ns |
| Muscle Force (a) vs Muscle Force (c) | 49200 | <0.0001 | *** |
| Muscle Force (b) vs Muscle Force (c) | 49200 | <0.0001 | *** |

*Kruskal-Wallis Test and Dunn's Multiple Comparison Test. $p < 0.05$ = significant, ns = not significant*

### Motoneuron Electrical Activity and Muscle Force Generation at 40 Hz Stimulation

At 40 Hz stimulation, significant differences were observed across all comparisons (p < 0.0001). The motoneuron with 5-HT receptors under physiological release (b) exhibited increased excitability compared to the no-receptor condition (a), whereas high 5-HT release (c) led to a decrease in excitability compared to both conditions (Table 1, Figure 2). Regarding the force curve, a shorter duration was observed in condition (b) compared to (a), though this difference was not statistically significant (p > 0.05). In contrast, the force curve in condition (c) was significantly shorter than in (a) and (b) (p < 0.0001). The force profile showed sustained activation for only about half the duration observed in the previous conditions, suggesting a fatigue effect (Table 2, Figure 2).

**Figure 2.**
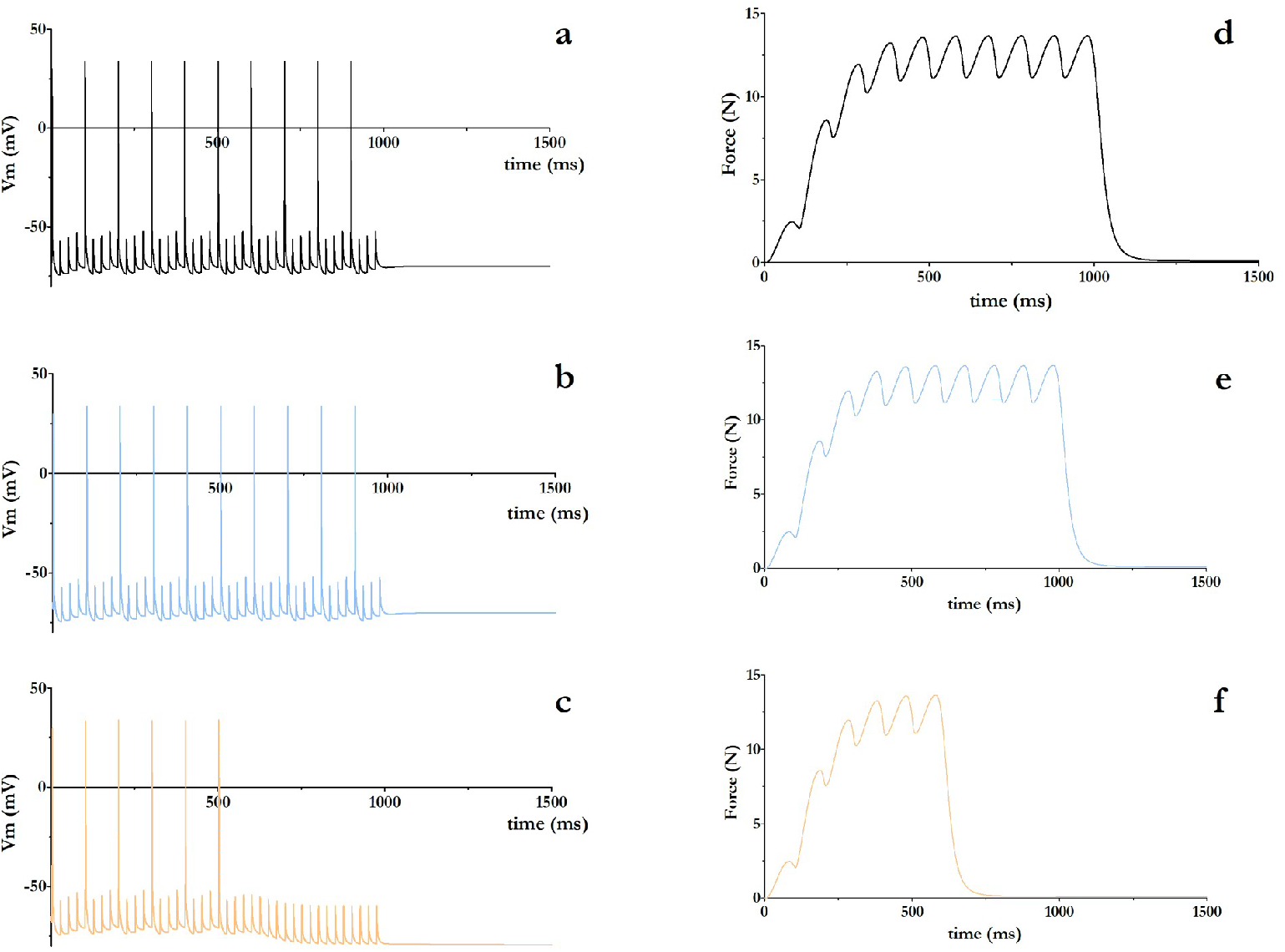
Membrane Potential of Motoneurons and Muscle Force Generated with 40 Hz Stimulation Frequency. Membrane potential of motoneurons without 5-HT receptors (a), with 5-HT1a and 5-HT2a receptors during physiological 5-HT release (b), and with 5-HT1a and 5-HT2a receptors during high 5-HT release (c). (d-f) Force generated by the muscle innervated by the corresponding motoneuron in (a), (b), and (c), respectively.

### Neuronal Activity and Muscle Force Generation with 100Hz Stimulation

At 100 Hz stimulation, significant differences were observed in all comparisons (p < 0.0001). The motoneuron with 5-HT receptors under physiological 5-HT release (b) showed an increase in excitability compared to the condition without receptors (a), whereas high 5-HT release (c) resulted in inhibition of the response compared to both previous conditions (Table 1, Figure 3). Regarding muscle force during 100Hz stimulation, the increased activation of 5-HT (c) generated a peak force similar to conditions (a) and (b), but with a significant decrease in the ability to maintain contraction over time (p < 0.0001). A “fatigue” effect was observed, where the initial force was high but rapidly declined (Table 2, Figure 3).

**Figure 3.**
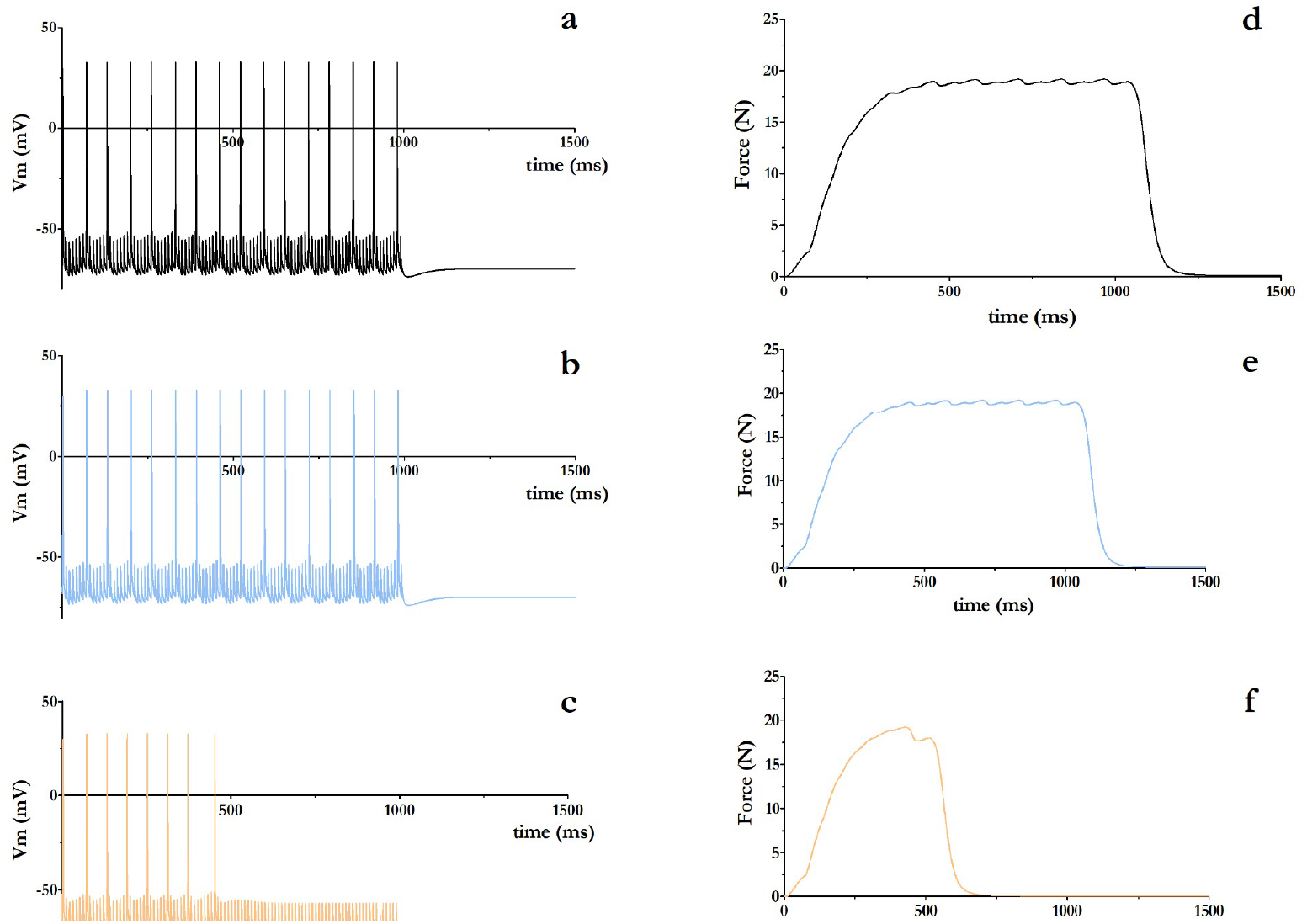
Membrane Potential of Motoneurons and Muscle Force Generated with 100 Hz Stimulation Frequency. Membrane potential of motoneurons without 5-HT receptors (a), with 5-HT1a and 5-HT2a receptors during physiological 5-HT release (b), and with 5-HT1a and 5-HT2a receptors during high 5-HT release (c). (d-f) Force generated by the muscle innervated by the corresponding motoneuron in (a), (b), and (c), respectively.

## Discussion

The results of this study demonstrate that the electrical activity of motoneurons and muscle force generation are significantly influenced by the presence of 5-HT receptors and the concentration of serotonin in the synaptic and adjacent environments. The implications of these findings are discussed below in the context of the existing literature and their relevance to motor control and certain neuromuscular pathologies.

### Serotonergic Modulation of Motoneuronal Excitability

Our results show that physiological 5-HT release increases motoneuron excitability, particularly at higher stimulation frequencies (40 Hz and 100 Hz). This effect is consistent with previous studies that have shown serotonin acts as an excitatory neuromodulator in motoneurons, facilitating action potential generation through the activation of 5-HT2a receptors [12, 13]. However, high concentrations of 5-HT resulted in a decrease in excitability, suggesting an inhibitory effect possibly mediated by excessive activation of 5-HT1a receptors, known for their role in neuronal hyperpolarization [8, 14].

### Implications for Muscle Force Generation and Fatigue

The generated muscle force showed a direct correlation with motoneuron activity. At a 10 Hz stimulation frequency, physiological 5-HT release did not significantly alter the force produced, suggesting that serotonin has no notable impact on muscle contraction at low-frequency stimuli. However, high 5-HT concentrations reduced contraction duration, indicating possible muscle fatigue induction. At higher stimulation frequencies (40 Hz and 100 Hz), physiological 5-HT release increased motoneuron excitability, which could facilitate a more sustained and powerful muscle response. However, exposure to high 5-HT concentrations resulted in a decreased ability to maintain contraction over time, evidencing a fatigue effect. This phenomenon could be explained by overstimulation of serotonergic receptors, leading to desensitization or disruption in motoneuron ionic homeostasis [4].

### Comparison with Experimental Studies and Model Validation

The findings from our computational model align with experimental studies investigating the role of serotonin in motor function. It has been reported that modulation of 5-HT receptors can influence muscle fatigue resistance, supporting the idea that a proper balance in serotonergic signaling is essential for maintaining motor function [3, 4, 15]. The consistency between our modeling results and existing experimental data suggests that the developed model is a valid tool for exploring the influence of serotonin on neuromuscular function. This model can serve as a foundation for future research aimed at understanding the underlying mechanisms of muscle fatigue and developing therapeutic interventions that modulate serotonergic signaling to improve muscle performance and recovery after injury. Furthermore, it allows the study of different ionic currents underlying neuromuscular excitability and inhibition phenomena.

### Study Limitations and Future Perspectives

Although our model provides valuable insights into the interaction between serotonin and neuromuscular function, it is important to acknowledge its limitations. Computational modeling simplifies the complexity of the neuromuscular system and may not capture all variables present in a living organism. Therefore, additional experimental studies are needed to validate and refine the model. Future research could focus on exploring interactions with other neurotransmitters, such as dopamine, adrenaline, and GABA, to understand how they affect motor function and fatigue resistance.

## Conclusions

The proposed model is consistent with the results of experimental studies. The findings suggest that fluctuations in serotonin concentration can affect muscle force generation, providing a model for studying fatigue induced by this neurotransmitter.

## List of Abbreviations

Not applicable.

## Author Contributions

JM and DP carried out the experiments. JM participated in the design of the study and performed the statistical analysis. JM and DP coordinated and helped to draft the manuscript. All authors read and approved the final manuscript.

## Ethics Statement

Not applicable.

## Data Availability Statement

Not applicable.

## Acknowledgments

We appreciate of academic support of Comisión Sectorial de Investigación Científica (CSIC) de la Universidad de la República, Uruguay.

